# Distinct depressive-like behavioural phenotypes in mice exhibit unique patterns of transcriptional perturbations across habenular cell subtypes

**DOI:** 10.64898/2025.12.23.696158

**Authors:** Priyam Narain, Marko Šušić, Aleksa Petković, Giuseppe Antonio-Saldi, Marc Arnoux, Mehar Sultana, Beemnet Andualem Belete, Dipesh Chaudhury

## Abstract

Major depressive disorder arises from diverse neurobiological mechanisms, yet how specific habenular cell types contribute to distinct depressive-like behaviours remains unclear. Using chronic social defeat stress, behavioural phenotyping, and single-cell RNA sequencing, we mapped stress-induced transcriptional changes across habenular cell types and lateral habenula (LHb) subregions. Our findings reveal that each behavioural phenotype is marked by its own transcriptional profile. These subregion-specific patterns across the LHb suggest that different cellular circuits contribute uniquely to depressive-like states.

## Main

The heterogeneity of major depressive disorder (MDD), which arises from distinct neurobiological substrates, poses significant challenges for treatment^1–5^. Current drug treatments are most often systemic, but they fail to target specific brain regions implicated in driving particular symptoms. Precision psychiatry is a promising approach for addressing this complexity through the identification and treatment of clinical subgroups that exhibit distinct neurobiological abnormalities^6–9^. By combining individual behavioural analysis along with molecular profiling, we can pinpoint cell-type and circuit-specific changes associated with MDD. This approach could reveal new drug targets inside unique brain regions for highly effective and precise treatments.

The habenula, a small epithalamic nucleus, has emerged as a critical brain region in the regulation of negative affect and aversion^10,11^. The habenula connects the limbic forebrain to the midbrain monoamine centers, such as the dorsal raphe nucleus (DRN) and the ventral tegmental area (VTA), both of which are involved in depressive-like behaviours following chronic stress^12–16^. Anatomically, the habenula is comprised of three distinct subnuclei: the medial habenula (MHb), lateral habenula (LHb), and perihabenula (PHb) - each with a different mosaic of neuronal subtypes^13,17–19^. The lateral habenula (LHb) plays a significant role in encoding negative reward prediction errors, punishment signals, and reward omission, with overall LHb hyperactivity suppressing reward-seeking^20–23^. LHb itself can be subdivided into four subregions: oval-medial, marginal, lateral, and subregion X (HbX).

Habenular hyperactivity has been strongly linked to mood disorders^16,24^ and is a therapeutic target for interventions such as ketamine administration and deep-brain stimulation^13,25,26^. In preclinical models, stress and other aversive experiences induce both hyperactivity and synaptic remodelling within the habenula^23,25,27–29^. This heightened activity shifts reward prediction errors toward negative values, contributing to depressive-like states^22^. Chronic stress further drives synaptic and transcriptional adaptations in the lateral habenula (LHb) that underlie these behavioural phenotypes^24,28,30^.

Hyperactivity of the LHb-VTA projection has been associated with passive coping^30^. Other studies found that LHb-DRN, but not LHb-VTA, is responsible for inducing the passive-coping phenotype^28,31^. Recently, activation of somatostatin-expressing cells in the HbX subregion of LHb was found to rescue passive-coping behaviour following chronic stress^32^. This opposing evidence suggests that finer cell-type resolution is required beyond that of projection-specificity to precisely map depressive-like behaviours to specific neuronal subpopulations. LHb displays substantial transcriptional heterogeneity, with at least four distinct transcriptionally and spatially delimited regions^19^. These subregions exhibit partial projection-specificity. In addition to habenular neuronal subpopulations, habenular astrocytes have been found to regulate both neuronal excitability and depressive-like behaviours^33–37^, providing a novel treatment target^38^ and suggesting an important role for habenular glia in the etiology of mood disorders. Despite growing evidence, our understanding of depression-associated cell-type-specific changes in the habenula remains limited.

Given the strong link between mood disorders and habenular dysfunction, and the lack of consistent evidence linking unique habenular projections with specific depressive phenotypes, a critical next step would be to investigate subregion-specific changes induced by chronic stress and relate these to distinct depressive-like phenotypes. To achieve this, we used single-cell RNA sequencing (scRNA-seq) to explore subregion-specific transcriptional changes in the Hb triggered by chronic social defeat stress (CSDS). CSDS is a translationally relevant murine model that induces a variety of stress phenotypes, unlike other stress paradigms^39–41^. Following CSDS, mice were classified according to three depressive-like traits: (1) social-avoidant (SI^+^), assessed by the social interaction test, (2) anhedonic (SPT^+^), assessed by the sucrose preference test, and (3) passive-coping phenotype (TST^+^), assessed by the tail-suspension test. Additionally, we identified (4) a resilient phenotype that did not exhibit depressive-like behaviour in any of the three behavioural assays, and (5) a susceptible phenotype that exhibited all three depressive-like traits.

## Results

### Behavioural results

Mice were exposed to 20 days of CSDS. A subset of stress-exposed animals (n = 42), along with stress-naive controls (n = 21), was assessed using the social interaction (SI), sucrose preference (SPT), and tail-suspension (TST) tests (Supp.Fig 1). To objectively define depressive-like traits, receiver operating characteristic (ROC) curves were generated using the social interaction ratio, sucrose preference percentage, and total immobility time as primary metrics^30^. Optimal cutoffs determined by the Youden J index were 104.2 (SI), 57.2% (SPT), and 203.8 s (TST) (Supp.Fig 2).

ROC analysis revealed better-than-random classification accuracy for SI and SPT (AUC = 0.61 and 0.56, respectively), validating the presence of social avoidance and anhedonia following CSDS. In contrast, the TST yielded a paradoxical reduction in immobility among stressed mice (AUC = 0.29), suggesting an increase in active, rather than passive, coping. For consistency, animals with the highest immobility duration that were SI⁻SPT⁻ were hand-designated as TST⁺. These defined behavioural subtypes provided the framework for subsequent transcriptional profiling of the habenula.

We selected five stress phenotypes for further analysis: SI⁺SPT⁻TST⁻ (SI⁺), SI⁻SPT⁺TST⁻ (SPT⁺), SI⁻SPT⁻TST⁺ (TST⁺), SI⁻SPT⁻TST⁻ (resilient), SI⁺SPT⁺TST⁺ (susceptible), in addition to stress-naive controls. As expected, nearly all control mice displayed the SI⁻SPT⁻TST⁻ phenotype.

### Identification of major cell classes

Using cell-type-specific marker genes, cross-referenced with a previously published dataset^19^, we identified 9 major cell clusters (Fig. 1A-B). Neurons were identified by expression of *Celf4* and *Snap25* and further classified into medial (MHb, 39809 cells) (Fig. 1C & D) and lateral habenula (LHb, 15505 cells) (Fig. 1E & F) by expression of *Tac2* and *Gap43*, respectively. Furthermore, *Mog*, *Olig1* and *Olig2* were used to identify oligodendrocytes (18831 cells), *Pdgfra* was used to identify polydendrocytes (1738 cells), *Col3a1* was used to identify fibroblasts (794 cells), *Slc6a11* and *Slc4a4* were used to identify astrocytes (4254 cells), and *Cldn5* was used to identify endothelial cells (6712 cells), *Abcc9* and *Pdgfrb* were used to identify pericytes (2075 cells). *Cx3cr1*, *C1qA*, *C1qB* and *C1qC* were used to identify microglia (9951 cells). *Mrc1* and *ApoE* were used to identify macrophages (868 cells). Several clusters that did not express any of the known marker genes remained unidentified, likely capturing unhealthy cells that were not removed by the initial filtering. These clusters were not considered for further analysis.

**Figure 1.**
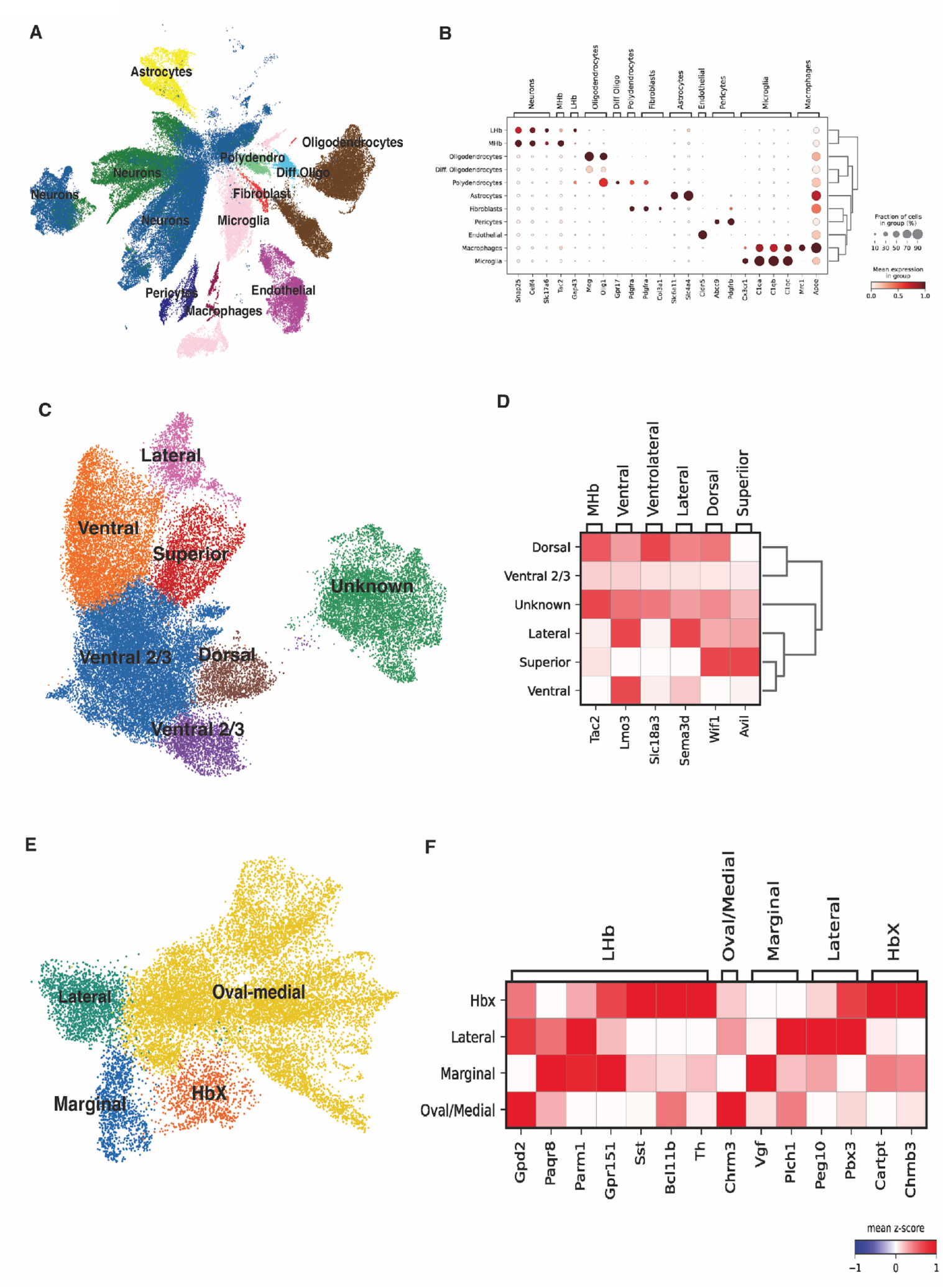
Single-cell transcriptomic profiling of mice habenula. (A)UMAP plot of all phenotypes integrated and processed, consisting of 98799 cells (MHb 39809, Oligodendrocytes 15653, LHb 15505, Microglia 9951, Endothelial 6712, Astrocytes 4254, Pericytes 207, Polydendrocytes 1738, Diff. Oligodendrocytes 1440, Macrophages 868, Fibroblasts 794) from n=22 animals. The clusters are annotated for a particular cell type based on canonical gene expression, with each cluster having the highest expression of a particular gene versus the other clusters. (B) Dot plot with dendrogram showing each cell class characterized by highly expressed marker genes with clusters annotated based on marker gene expression. The color of each box indicates the mean expression of each gene in a particular cluster, with red indicating the highest and blue lowest mean expression.(C &E) UMAP plot of MHb and LHb neurons, respectively. (D & F). Expression of MHb and LHb subtype-specific marker genes across neuronal subtypes.

### Phenotype-specific patterns of differential expression across major cell classes

We first assessed the cell-type composition within each distinct cell population, ensuring sufficient cellular representation for subsequent analyses (Supp.Fig. 3). To identify associations between specific phenotypes and cell types in the habenula, we performed differential expression (DE) testing using the Wilcoxon rank sum test to identify phenotype-specific transcriptional changes across major cell types with stress-naive mice utilized as the comparison baseline. With a Bonferroni-adjusted *p* value ≤ 0.05 and absolute log_2_ fold change (LFC) ≥0.25, we identified a large number of differentially expressed genes (DEGs) in the whole habenula for each stress-induced phenotype relative to stress-naive mice. In total, TST^+^ mice displayed the highest number of DEGs (14256), followed by resilient (12325), susceptible (7574), SI^+^ (2367), and SPT^+^ (1985) mice. Behavioural phenotypes widely differed in the proportion of total DE that is attributable to a specific cell type (Fig. 2A). To further establish association between phenotypes and cell types, we identified unique and shared DEGs across phenotypes for each major cell type (Fig. 2A and Supp. Fig. 4). The most unique transcriptomic signature with relevant number of DEGs was found in the LHb neurons and was associated with the susceptible phenotype. Amongst the glia, the oligodendrocytes had the highest number of DEGs (6025) found in the resilient phenotype.

**Figure 2.**
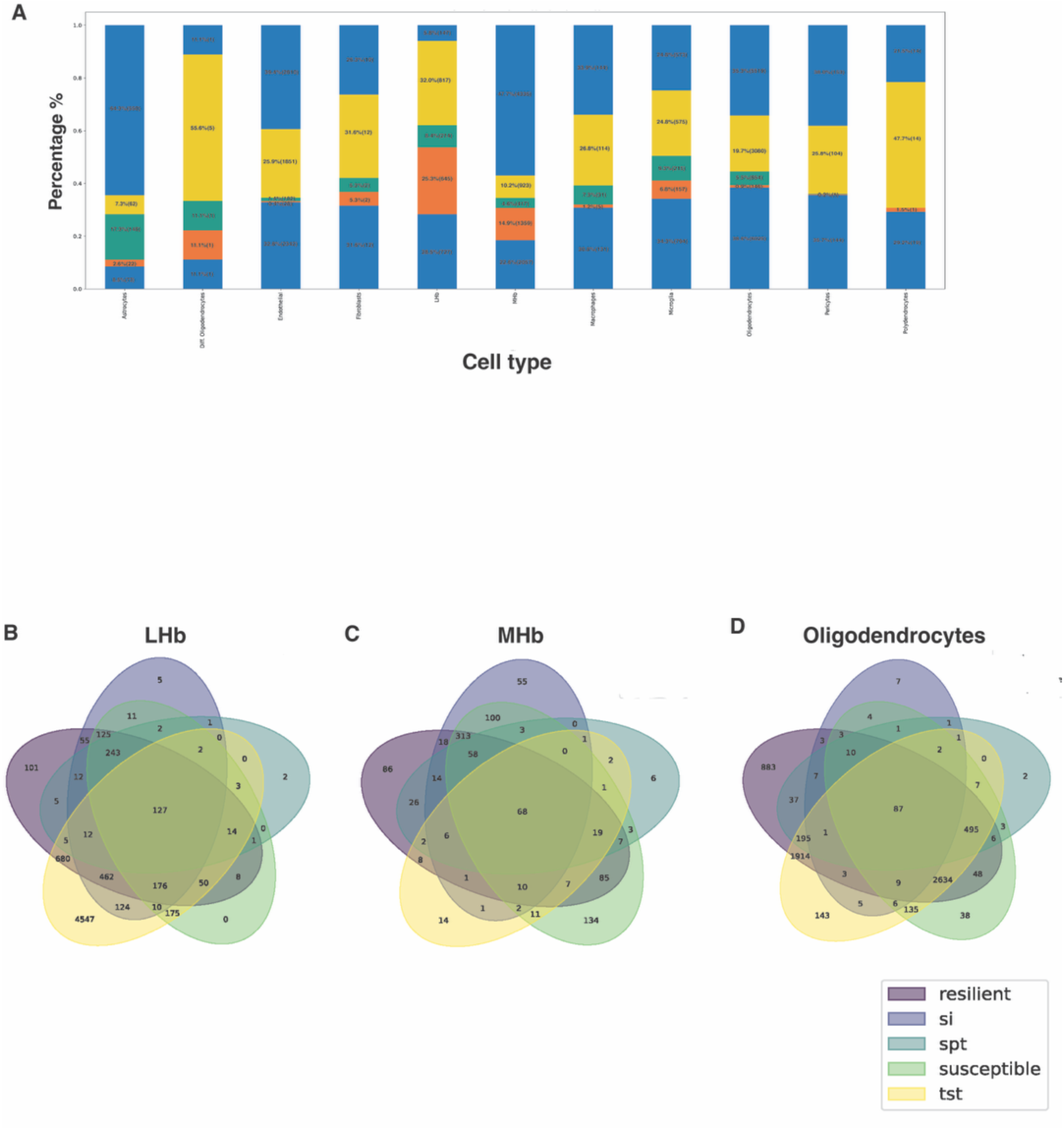
Cell type-specific distribution and transcriptional overlap of differentially expressed genes (DEGs). (A) Stacked bar plot representing the proportional distribution of differentially expressed genes across distinct behavioural phenotypes, including TST+, Susceptible+, SI+, SPT+, and Resilient groups. Each bar denotes a specific cell type, and the y-axis indicates the percentage of DE with each phenotype. Cell types include Oligodendrocytes, Astrocytes, Microglia, Medial Habenula (MHb) neurons, Lateral Habenula (LHb) neurons, Pericytes, Polydendrocytes, Endothelial cells, and Macrophages. (B–D) Venn diagrams illustrating the overlap of DEGs identified in phenotypic comparisons relative to control animals within three major cell populations: (B) LHb neurons, (C) MHb neurons,(D) Oligodendrocytes. Each diagram displays the number of unique and shared DEGs among Resilient, SI+, SPT+, TST+, and Susceptible groups.

### Identification of LHb subregions

Since LHb has specifically been associated with mood disorders, we subsetted LHb neurons and further distinguished subregions. We used region-specific LHb neuronal markers, cross-referenced with a previously published dataset (Wallace et al., 2020), to identify 4 subregions of LHb (Fig. 1E-F). Briefly, *Chrm3* was used to identify the oval-medial subtype (11464 cells) *Gap43* and *Vgf* were used to identify the marginal subtype (1873 cells), and *Gpr151* and *Cartpt* were used to identify lateral (1020 cells) and HbX (1148 cells) subregions of LHb, respectively.

### Phenotype-associated patterns of differential gene expression across LHb subregions

We performed DE testing between the stress-induced phenotypes and stress-naive mice as before for each LHb subregion (Fig. 3A). Distribution of DEGs across the LHb subregions for the five selected phenotypes (Fig. 3B-F) revealed a consistent pattern where the oval-medial subregion had the largest proportion of LHb DEGs for resilient (69.2%, DEGs = 1121), SI^+^ (88.7%, 1594), SPT^+^ (93.5%, 501), and susceptible (85.5%,1298) phenotypes, suggesting its general non-phenotype-specific involvement in post-stress transcriptional response. In contrast, the HbX region demonstrated a robust association with the TST^+^ phenotype. This subregion exhibited a highly distinct and specific transcriptomic signature, contributing the majority of LHb DEGs (85.9%, 2588) in the TST^+^ mice. This supports the regional specificity of stress-induced DE, particularly the importance of the oval-medial subregion for most stress phenotypes and the unique involvement of HbX in the TST^+^ phenotype.

**Figure 3.**
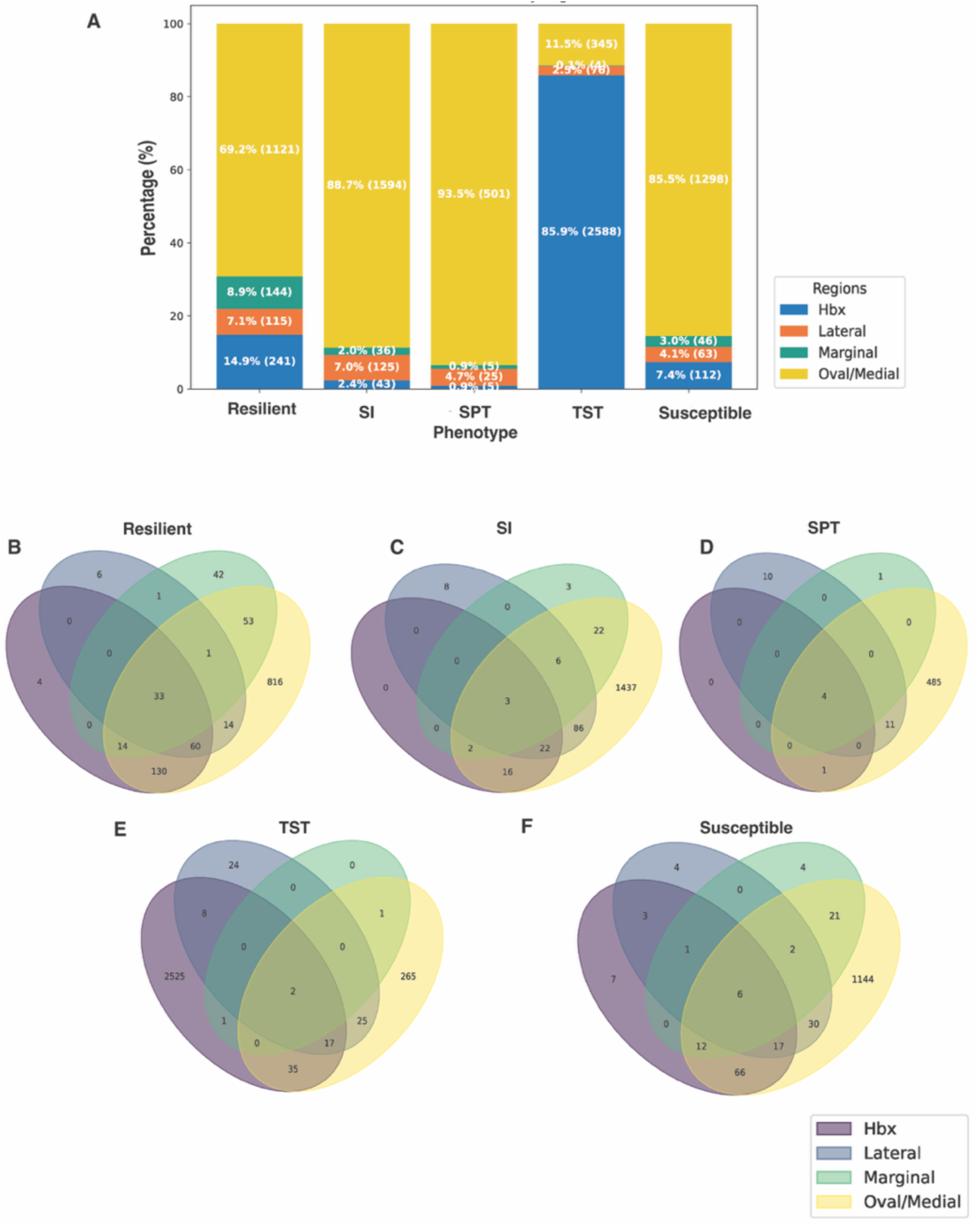
DEGs across lateral habenular cell types and behavioural phenotypes. **(A)** This stacked bar graph displays the distribution of differentially expressed genes (DEGs) across different sub-regions of the lateral habenula (Hbx, Lateral, Marginal, Oval/Medial) for different phenotypes (Resilient, SI, SPT, Susceptible, TST). The y-axis represents the percentage of DEGs associated with each region, while the x-axis denotes the behavioural phenotypes. The absolute number of DEGs for each region within each condition is provided in parentheses next to its percentage. This visualization aims to associate specific regional gene expression changes with particular behavioral or physiological phenotypes.Overlap and specificity of differentially expressed genes across LHb sub-regions in different phenotypes. **(B-F)** Venn diagrams illustrating the overlap and unique distribution of DEGs across four distinct sub-regions of the LHb: Hbx (purple), Lateral (blue), Marginal (green), and Oval/Medial (yellow). B: Resilient: DE genes Phenotype vs Control C: SI: DE genes Phenotype vs Control D: SPT: DE genes Phenotype vs Control E: Susceptible: DE genes Phenotype vs Control F: TST: DE genes Phenotype vs Control

We further analyzed the direction of gene regulation (upregulated vs. downregulated) and compared the unique numbers of DEGs in each subregion, as described below. Due to strong and specific responses, we further examined pathway enrichment patterns in the oval-medial and HbX between the phenotypes using Ingenuity Pathway analysis (IPA).

### *Oval-medial* (Fig. 4A)

In the oval-medial subregion, SI^+^ mice had the highest number of DEGs (Total = 1594, Up = 45, Down =1549), followed by susceptible (1298, 100, 1198), resilient (1121, 82, 1039), SPT^+^ mice ( 501, 21, 480) and TST^+^ mice (345, 76, 269). Here, SI^+^ and susceptible mice exhibited the most extensive gene expression changes. These two behavioural phenotypes share a substantial number of downregulated genes, suggesting a common underlying molecular response. Resilient mice show an intermediate degree of gene alteration, while SPT^+^ and TST^+^ mice display the least perturbed gene expression in this subtype.

**Figure 4.**
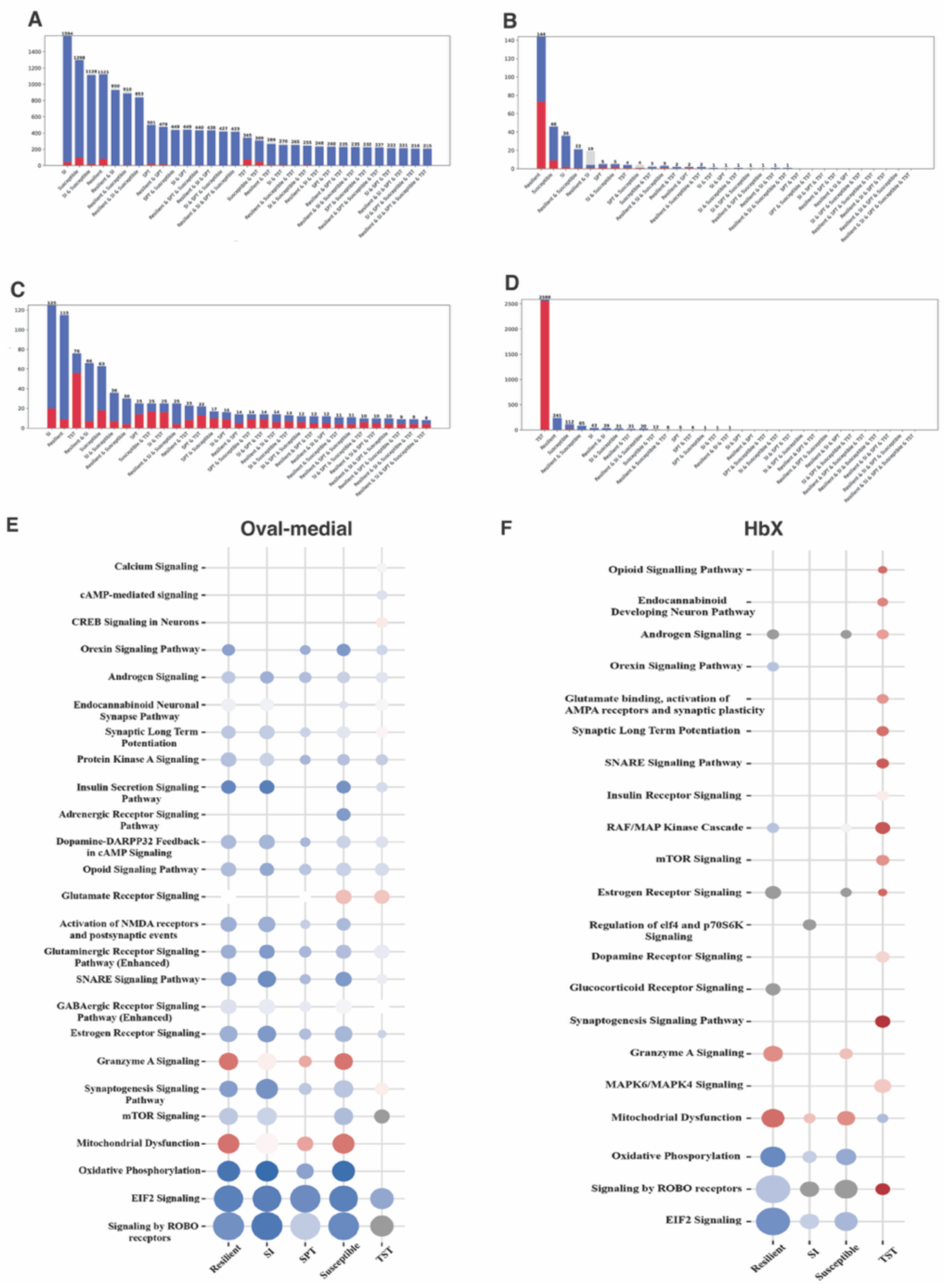
Common and distinct transcriptional responses across different behavioural phenotypes. **(A - D)** DEG distribution and overlap by phenotype. The y-axis represents the number of DEGs, while the x-axis shows individual or combined phenotypic comparisons. DEGs are color-coded by direction: red for upregulated genes, blue for downregulated genes, and gray for genes with mixed or conflicting regulation.(A) Oval medial (B) Marginal (C)Lateral (D)HbX **Bottom panel : (E & F)**Pattern of pathway enrichment across the phenotypes provided by Qiagen Ingenuity Pathway Analysis (IPA) software. Warmer colors (positive z-scores) indicate pathway activation, while cooler colors (negative z-scores) indicate pathway inhibition. Dot size indicates the negative log10 of the p-value with larger dots corresponding to more significantly enriched pathways.

To distinguish unique molecular signatures of resiliency and susceptible phenotype, we identified unique DEGs in both. Resilient and susceptible animals exhibited 211 and 388 unique DEGs, respectively. In the resilient animals, *Kcnj10* was significantly downregulated (log_2_FC = –1.50), while *Cacna1b* was upregulated (+0.78), potentially increasing hyperexcitability in these cells. In the susceptible animals, several genes related to synaptic function and excitability were upregulated, including *Grid2* (+0.24), *Grin2b* (+1.27), *Cacna2d3* (+1.28), and *Kcnc2* (+1.04), which likely bias these cells towards high-frequency firing and overall rapid sustained excitatory signalling.

To further explore differences between the phenotypes in this subregion, we performed IPA pathway enrichment analysis. Resilient animals showed significant predicted inhibition in ’Oxidative phosphorylation’ (-log(B-H p-value) = 30.1, z = -6.403), ’EIF2 signalling’ (61, -5.778), and ’ROBO signalling’ (71.8, -5.014) pathways. These pathways also appeared in the SI^+^ and SPT^+^ groups with slightly lower enrichment values. ’Synaptogenesis signalling pathway’, ’mTOR signalling’, and ’Insulin secretion signalling pathway’ have a similar inhibition pattern in resilient, SI^+^, and Susceptible groups. In contrast, ’Granzyme A signalling’ and ’Mitochondrial dysfunction’ showed positive activation-associated z-scores in the resilient, SPT^+^, and susceptible phenotypes. The TST^+^ group showed limited enrichment overall, with relatively low significance and minimal activation, except for ’EIF2 signalling’ (32.6, -3.873), which remained highly enriched. Notably, “Calcium signalling” (2.59, -0.33), “CREB signalling” (4.01, 0.655), and “cAMP signalling” (2.74, -1.265) appeared only in TST^+^.

### *Marginal* (Fig. 4B)

In the marginal subregion, resilient mice had the highest number of DEGs (Total = 144, Up = 73, Down = 71), followed by susceptible (46, 9, 37), SI^+^ (36, 2, 34), SPT^+^ mice (5, 2, 3), and TST^+^ mice (5, 1, 3). Overall, these DE patterns suggest low relevance of this subregion for the emergence of stress-induced phenotypes.

### *Lateral* (Fig. 4C)

In the lateral subregion, SI^+^ mice had the highest number of DEGs (n=125, Up=20, Down=105), followed by resilient (115, 9, 106), TST^+^ (76, 55, 21), and susceptible mice (66, 15, 51). Hence, no clear phenotypic pattern emerged.

### *HbX* (Fig. 4D)

In the HbX subregion, TST^+^ mice had the highest number of DEGs (n=2588, Up =2561, Down = 27), followed by resilient (241, 6, 235) and susceptible (112, 21, 91). These patterns suggest that the HbX region is uniquely tied to the presence of the TST^+^ phenotype or buffering against it. In addition, resilient and susceptible animals exhibited 156 and 27 unique DEGs, respectively.

From our IPA analysis, the TST^+^ group displayed the broadest and most distinct pathway profile compared to susceptible, resilient, SI^+,^ and SPT^+^ groups. Specifically, TST^+^ animals showed activation of signalling pathways, including ’Opioid signalling’ ( -log(B-H p-value)= 1.31, z = 4.629), ’Glutamate binding and AMPA receptor activation’ (2.07, 3.317), ’Synaptic long-term potentiation’ (2.35, 4.536), and the ’SNARE signalling pathway’ (2.43, 5.209). ’Synaptogenesis’, ’mTOR signalling’, and ’MAPK6/MAPK4 signalling’ appeared exclusively in TST^+^. Overall, the TST^+^ group showed the broadest pathway profile in the HBx region, driven by synaptic and intracellular signalling.

## Discussion

Our findings demonstrate that CSDS induces diverse and complex behavioural phenotypes in mice, extending beyond a simple susceptible/resilient dichotomy based on social avoidance^42^. Where possible, we used unbiased thresholds from ROC curves to construct composite behavioural phenotypes and investigate molecular underpinnings of the associated habenular stress responses. We identified 9 major cell clusters, including distinct MHb and LHb neuronal populations, using previously established gene markers^19^. Our findings on the transcriptional heterogeneity align well with recent scRNA-seq studies that have comprehensively defined cell populations and their spatial organisation in the mammalian habenula^18,19^.

We were unable to obtain a reliable ROC classifier for the immobility time in the TST. We interpret this finding as evidence for a distinct behavioural adaptation driven by the specific nature of the stressor. Rather than inducing passive coping, the CSDS paradigm promotes an active, or proactive, coping strategy. Chronic stress paradigms that are uncontrollable and unpredictable, such as UCMS, are known to promote passive coping responses (i.e., increased immobility), perhaps reflecting a state of helplessness^41^. The CSDS paradigm, however, is an ethologically distinct stressor involving direct, agonistic encounters. In this context, active behaviours like struggling or attempting to escape may be reinforced as a means to terminate or mitigate the stressful social encounter. The observed decrease in immobility could therefore be interpreted not as an absence of a stress response, but rather a signature of an adaptive, active coping phenotype. Notably, our finding that CSDS exposure leads to decreased immobility in the TST, indicative of an active coping strategy, stands in contrast to studies using other chronic stress paradigms. For instance, Cerniauskas and colleagues demonstrated that mice subjected to UCMS display a more canonical phenotype of increased TST immobility duration^30^. This divergence strongly suggests that chronic stress does not produce a single, universal behavioural outcome. Instead, the consequences are critically shaped by the specific nature of the stressor, with social defeat potentially favouring proactive, rather than passive, coping mechanisms. Therefore, instead of using ROC classification, we designated animals with the highest immobility scores as TST^+^. Interestingly, this approach disclosed a strong association between the TST^+^ group and the HbX cell type. In light of our ROC analysis, which suggests that CSDS promotes active coping behavioural strategies, the HbX association likely represents a molecular signature of passive coping, highlighting that molecular signatures could be associated with specific behavioural strategies.

Glia-neuron interaction in the LHb has been shown to modulate neuronal firing. For example, astroglial potassium channels (Kir1.4) were shown to be upregulated in mice exposed to CMS, resulting in increased burst firing of LHb neurons^34^. Our observations that astrocytes and oligodendrocytes also show different stress-induced changes in their transcriptomic profiles may be indicative of some of the differences in the functional coding of various behavioural phenotypes. Moreover, it also adds to the novel role of glia-neuron interaction in the context of differential stress responses. We also show an association of DE in oligodendrocytes with the resilient phenotype. Oligodendrocytes are vital for neuronal support through myelination. New myelin formation in the medial prefrontal cortex (mPFC) of resilient mice has been shown to be an adaptive response to repeated aggression, facilitating neuronal circuits that mitigate the negative effects of stress^43^. Conversely, emerging evidence suggests that oligodendrogenesis and myelination in the PFC are highly sensitive to stress. Prolonged social isolation can induce transcriptional and structural changes in PFC oligodendrocytes, which impair myelin formation^44^. Our association of resiliency with oligodendrocyte DE adds to existing reports of differential plasticity of oligodendrocytes and progenitor cells across stress-associated brain regions^45^. Furthermore, it is known that oligodendrocytes can enhance neuronal firing through both improved myelination and TRKB-dependent modulation of presynaptic release, suggesting a glial mechanism for strengthening stress-relevant circuits. This aligns with evidence that resilient mice preserve coherent reward-related firing in BLA circuits after social defeat^46^, and BNST CRF neurons show stress-history dependent increases in excitability that support resilient behaviour^47^. Likewise, human imaging indicates that resilient coping involves dynamic up-regulation of prefrontal and insular activity under stress^48^. Taken together, this evidence suggests that oligodendrocyte-supported firing may help drive resilient circuit activity.

We also observed a strong association of DE in LHb neurons with the susceptible phenotype, which is consistent with a large body of evidence regarding hyperexcitability of habenular neurons in depressive-like mice ^24,29,49–52^. Following CSDS, LHb neurons projecting to the dorsal raphe nucleus (DRN) in susceptible mice exhibit daytime-specific elevated firing relative to resilient and stress-naive mice^53^. Another study found that, following chronic mild stress, the LHb cells projecting to the ventral tegmental area (VTA) rather than the DRN exhibit elevated firing^30^, highlighting the importance of habenular long-range projections as well as the varying post-stress effects of different chronic stress paradigms^41^. A more recent study examined the activity of LHb cells during CSDS and found that their elevated activity from the first day of the paradigm biases the animals towards the SI^+^ phenotype^24^.

Enhanced serotonergic signalling from the dorsal raphe projections to the LHb promotes resilience to stress^54^. Moreover, dense 5HT terminals innervate lateral and medial zones of the LHb^55^. Serotonergic G-protein-coupled receptors that can activate (via Gq or Gs G-proteins) or inhibit (via Gi G-proteins) neural activity. The distribution of 5-HT receptor subtypes in subregions of the LHb is one likely mechanism by which different responses to stress correlate to particular gene expression profiles, since different subtypes of G-proteins have different effects on the intracellular signalling cascade. The expression analysis across four distinct anatomical LHb subregions —oval-medial, marginal, lateral, and HbX—revealed a highly heterogeneous and phenotype-specific molecular landscape in response to chronic social stress. Unique DEGs and direction of regulation were associated with specific subregions and phenotypes. The oval-medial subregion shows overall downregulation, particularly in SI^+^ and susceptible mice. Furthermore, a shared molecular signature associated with stress exposure was evident, as the largest number of downregulated genes were common to the resilient, SI^+^, and susceptible phenotypes, with a significant overlap also observed between SI^+^ and susceptible animals. Interestingly, resilient mice displayed the highest number of uniquely upregulated genes, followed by the susceptible and SI^+^ groups (Supplementary fig 5), potentially reflecting adaptive or maladaptive molecular responses. These findings align with earlier research demonstrating functional differentiation within the LHb, where stressors preferentially activate the medial portion^56,57^. Stress increases the activity of excitatory projections from the entopeduncular nucleus (EP) to the oval-medial subregion of the LHb^30^. Furthermore, stress-induced increased glutamatergic signalling from EP to the oval-medial subregion of the LHb likely activates calcium signalling pathways that lead to changes in gene expression profile in different behavioural phenotypes.

Other subregions had more balanced patterns of up- and downregulation. Lateral and marginal subregions showed overall smaller, more balanced, and more uniform responses across the phenotypes. Furthermore, in contrast to the oval-medial subregion, where all phenotypes showed significant differential expression, HbX exhibited extensive gene upregulation specifically in the TST^+^ phenotype. In other phenotypes, HbX shows either minimal changes or predominantly downregulation. Neuropeptide CCK signalling into the HbX has been implicated in encoding stress response following exposure to CMS, where decreased firing in somatostatin-expressing neurons was associated with anhedonic response in SPT^32^. The same study found that activation of these cells in the HbX subregion rescued the passive coping phenotype. Thus, it is possible that CCK signalling into the HBx may be responsible for the specific transcriptomic profile we observed for TST^+^ mice. In contrast, the SPT^+^ phenotype in our study presented a unique gene signature exclusively within the oval-medial subpopulation, but not elsewhere. This difference may be related to molecular responses to different stressors. Our results suggest the general importance of the oval-medial subregion for post-stress phenotype modulation and the specific importance of the HbX subregion for the TST^+^ phenotype.

## Conclusion

In summary, our study highlights the heterogeneous nature of cellular chronic stress response within the habenula. By investigating transcriptomic responses in the anatomically distinct subregions of the LHb, a key brain region implicated in depression, we have identified specific transcriptomic profiles associated with different depressive-like phenotypes, potentially contributing to stress-induced phenotypes like social avoidance, anhedonia, and passive coping. Understanding cell-type-specific molecular mechanisms underlying diverse stress-triggered behavioural responses is key to fully unravelling the intricate molecular underpinnings of depressive symptomatology.

## Contributions

P.N., M.Š., A.P. and D.C. conceptualized the study. A.P. developed and, together with M.Š., carried out the behavioural segment of the study. M.Š. analysed the behavioural data. P.N. and M.Š. ran scRNA sequencing with M.A. and M.S assisting in the process. G.A.S., M.Š., and P.N. analysed the transcriptomics data. B.A. performed the enrichment analysis. P.N. and A.P. interpreted the data. P.N, A.P. and D.C. wrote the manuscript. P.N. and D.C. supervised and secured funding.

## Materials and methods

### Animals and ethics statement

Adult wild-type C57BL/6J mice (>7 weeks old) were imported from Jackson Laboratory (Bar Harbor, MS, USA). Retired CD1 breeder mice from Charles River laboratory (Wilmington, MS, USA) were used as resident aggressors. All experimental mice were maintained within standard controlled housing conditions at a temperature of 21±2°C, humidity of 50±10%, with *ad libitum* access to food and water and enrichment in the form of wood shavings. Mice were kept on a 12-12 light/dark cycle (light onset at 7am - ZT0, light offset at 7pm – ZT12). Animal protocols have been approved by the National Institute of Health Guide for Care and Use of Laboratory Animals (IACUC Protocols: 150005A2, 19-0004A1) and NYUAD Animal Care and Use Committee.

### Behavioural paradigm

A cohort of C5BL/6J mice (n = 71) was exposed to 20 days of chronic social defeat stress (CSDS). A subset of the experimental cohort (n=42), along with a control group (21 mice), was examined for depressive-like phenotype using three standard behavioural assays: social interaction test (SI), sucrose preference test (SPT), and tail suspension test (TST). The SI test and TST tests were conducted approximately 24 and 48 hours after the last defeat session, while SPT was run in parallel over 4 days (Supp.Fig.1).

### Chronic social defeat stress (CSDS)

The chronic social defeat stress (CSDS) paradigm was performed as described previously with minor modifications^42,53,58^. Briefly, the C57BL/6J experimental mice were exposed daily to 10 minutes of direct contact with a novel CD1 mouse (physical stress) followed by 24 hours of sensory contact. The procedure was carried out daily for 21 days between ZT8 and ZT10. After the conclusion of the CSDS, all animals were single-housed for 24 hours before the start of behavioural testing.

### Social interaction (SI) test

Social avoidance behaviour in C57BL/6J experimental mice towards a novel non-aggressive CD1 mouse was assessed using a two-stage social interaction (SI) test^42^. Each stage lasted for 180 seconds. In the first stage, the experimental mouse was allowed to freely explore a square-shaped arena (44 × 44 cm) containing an empty perforated Plexiglas mesh (10 × 6 cm). The area (14 x 26 cm) around the center of the mesh was defined as the social interaction zone. In the second stage, a novel non-aggressive CD1 social target was introduced into the perforated mesh. Behavioural recording was performed using the TopScan video tracking system (CleverSys Inc.). The social interaction (SI) score was calculated as [100 × (time spent in interaction zone with social target present) / (time spent in the interaction zone with social target absent)]. SI-positive (avoidant) and SI-negative mice were segregated based on a threshold obtained from a receiver operating characteristic (ROC) curve constructed on all of the obtained SI scores. The Youden J index was used to determine the optimum unbiased cutoff threshold separating SI^+^ and SI^-^ mice^59^.

### Tail suspension test (TST)

Passive coping behaviour in C57BL/6J experimental mice was measured using the tail suspension test (TST) as described previously^57^. The experimental C57BL/6J mice were suspended by the tail 40 cm above the ground for 6 minutes. Each trial was recorded using a fixed Nikon 6000 camera and manually scored in a double-blind procedure. An ROC curve constructed from the total immobility time and the Youden J index was used to determine the optimum unbiased cutoff threshold separating TST^+^ (passive-coping) and TST^-^ mice. Since this approach did not yield a good classifying threshold, we chose the animals with the longest immobility time in the cohort as TST^+^.

### Sucrose preference test (SPT)

Anhedonic behaviour in mice was measured using the sucrose preference test (SPT) as described previously^60^. After completion of the SI test, all mice were single-housed and habituated to two liquid diet feeding tubes (Bio-Serv.) containing water for 2 days. Following the tube habituation, mice were water-deprived for 24 hours. Starting at ZT12, the animals were exposed to one tube containing 40 ml of water and another tube containing 40 ml of 1% sucrose solution. Each tube was weighed before the start of the test. Bottles were reweighted and switched in position at ZT2, ZT8, and ZT11 on the following day. Following the test conclusion, regular water bottles were reintroduced. Liquid consumption was calculated for each consecutive interval as well as for the total 24-hour period. Sucrose preference (SP) score was calculated as [100% × (amount of sucrose consumption) / (amount of sucrose consumption + amount of water consumption)]. An ROC curve constructed from the SP scores and the corresponding Youden J index was used to determine the optimum unbiased cutoff threshold separating SPT^+^ (anhedonic) and SPT^-^ mice.

### ROC curve and behavioural categorization

The data collected from these behavioural assays showed substantial variability in individual test results for both control and experimental mice, making it difficult to assess the effects of CSDS on individual animals. To address this, we constructed receiver operating characteristic (ROC) curves, an objective method commonly used in clinical research to evaluate binary classifiers. For the ROC curves, we used the social interaction ratio as the primary measure for the SI test, the sucrose preference percentage for the SPT, and the total immobility time for the TST. These measures were used to estimate optimal threshold values based on the Youden J index, which helped us determine whether each animal exhibited depressive-like or control-like behaviour. The optimal cutoff values, representing the highest true-positive rates and lowest false-positive rates, were 104.2 for the SI test, 57.2% for the SPT, and 203.8 seconds for the TST. However, we failed to establish a reliable objective threshold for the TST through this method. Hence, we deemed as TST^+^ the animals that exhibited the longest immobility times and were also SI⁻ and SPT⁻ (i.e., not overlapping with the other behavioural phenotypes). Susceptible animals were SI^+^ and SPT^+^ with the longest immobility times.

### Single-cell preparation and cDNA library construction

Single-cell RNA sequencing (scRNA-Seq) was performed on the habenular nuclei of selected mice (Supp.Fig 1). Briefly, mice were perfused for 40-60 seconds with 20 mL of ice-cold artificial cerebrospinal fluid (aCSF) containing: 128 mM NaCl, 3 mM KCl, 1.25 mM NaH_2_PO_4_, 10 mM D-glucose, 24 mM NaHCO_3_, 2 mM CaCl_2_, and 2 mM MgCl_2_. The aCSF was continually oxygenated with a mixture of 95% O_2_ and 5% CO_2_. Brain slices (250 μm in thickness) containing the Hb were cut using a microslicer (DTK-1000, Ted Pella) in ice-cold sucrose aCSF. Slices containing the habenular complex were examined immediately under a stereoscope and 0.5μm diameter habenular tissue punches were collected into papain dissociation buffer comprising of 10U/ml Papain (Worthington dissociation system, LK003150), 100 U/ml DNase1 (Worthington dissociation system, LK003150), 1X Glutamax (Gibco), 0.2X B27 supplement (Thermo Fisher Biosciences), and 1% w/v D(+)trehalose (Sigma) in hibernate A medium (Life Technologies). The collected tissue samples were incubated for 45 minutes at 37 °C and triturated to facilitate cell dissociation. The enzymatic digestion was stopped by 10 mg/ml ovomucoid protease inhibitor with bovine serum albumin (Worthington dissociation system, LK003150), and samples were kept chilled for the rest of the dissociation procedure. The final suspension was obtained in 0.04% BSA and was then filtered through a 40μm pluriStrainer filter (pluriSelect). The resulting suspension was loaded into a 10X Genomics Chromium single-cell chip at a concentration of ∼400 cells/μL. Downstream sequencing library preparation was performed using the 10X Genomics Chromium Single Cell Kit V3.1 according to the manufacturer’s guidelines. The libraries were sequenced on an Illumina NovaSeq 6000 instrument using the instructions provided by 10X Genomics.

### Single-cell RNA-seq data preprocessing and quality control

Raw gene expression matrices were loaded into an AnnData object using the Scanpy Python package^61^. Gene and cell identifiers were first made unique. A batch identifier was added. Initial quality control (QC) was performed to remove low-quality cells and genes. First, we identified mitochondrial genes by matching gene names with the prefix ^(mt)-. QC metrics, including the number of genes per cell, total counts per cell, and the percentage of counts from mitochondrial genes, were calculated. Genes detected in fewer than 3 cells and cells expressing fewer than 200 genes were removed from the dataset. Cells with outlier QC metrics were flagged but retained for analysis; these included cells with total counts below the 5th or above the 95th percentile, and cells where the mitochondrial gene content exceeded the 95th percentile.

### Doublet Detection

Potential doublets in each single-cell dataset were identified and removed using scvi-tools. First, a scVI model was initialised and trained on the raw count matrix of the dataset to learn a harmonised latent representation of the cells. This pre-trained scVI model was then used to initialise the SOLO doublet detection model. After training the SOLO model, it was used to predict a doublet probability score for each cell. A final, hard classification of either ’singlet’ or ’doublet’ was assigned to each cell, and these annotations were used for subsequent filtering.

### Integration, Normalisation, Scaling, Feature Selection and Clustering

The 18 datasets were concatenated with sc.concat, adding dataset id to .obs[’dataset’], the raw count matrix (.layers[’counts’]) was used as the primary input for analysis. Cell counts were first normalised by library size to a target sum of 10,000 counts per cell, followed by a natural log-transformation (log(X+1)). The data was then scaled to unit variance and zero mean.

Principal Component Analysis (PCA) was performed on the scaled expression matrix to generate an initial 50-dimensional embedding. To correct for batch effects originating from different datasets (as defined in .obs[’dataset’]), we applied the Harmony integration algorithm to this PCA embedding, with a maximum of 50 iterations. Finally, a k-nearest neighbour (k-NN) graph was constructed on the resulting Harmony-corrected latent space (X_pca_harmony) to be used for downstream clustering and visualisation. A nearest-neighbour graph was constructed based on the Euclidean distances in the X_pca_harmony space. This graph was used to compute a two-dimensional embedding for visualisation using Uniform Manifold Approximation and Projection (UMAP). Finally, cell communities were identified by applying the Leiden clustering algorithm to the nearest-neighbour graph with a resolution parameter of 0.4.

### Cell-type identification and marker gene analysis

An AnnData object containing the lateral habenula (LHb) single-cell data was loaded for analysis. Cell clusters were defined based on the Leiden clustering algorithm and were subsequently annotated into distinct cell types, including ’Oval/Medial’, ’Hbx’, ’Marginal’, and ’Lateral’ subtypes. This annotation was based on the expression of known marker genes. A matrix plot was generated to visualise the mean z-score of log-normalised expression (l1p layer) for these marker genes across the identified cell types.

### Differential gene expression analysis

To identify differentially expressed genes (DEGs) between different experimental phenotypes (’resilient’, ’susceptible’, etc.) and the control group, we performed a Wilcoxon rank-sum test using Scanpy’s rank_genes_groups function. This analysis was conducted in two ways:

1. Across the entire LHb dataset to identify global changes.
2. Within each annotated sub-region (’Oval/Medial’, ’Hbx’, etc.) to identify region-specific changes.

For these analyses, raw counts from the ’counts’ layer were used, which were then normalized to a target sum of 10,000 and log-transformed. Genes were considered differentially expressed if the adjusted p-value was less than or equal to 0.05. The absolute log-fold change was also calculated for further filtering and interpretation.

### Bootstrap analysis for identification of stable DEGs

To ensure the robustness of our DEG findings, we implemented a bootstrap analysis with 5,000 iterations for each cell type within the dataset. In each iteration, cells were downsampled to an equal number per condition to avoid biases from varying cell numbers. The Wilcoxon rank-sum test was performed on each bootstrap sample to identify DEGs.A gene was considered a ’stable’ DEG if it was found to be significant (adjusted p-value ≤ 0.05) in at least 50% of the bootstrap iterations. The results were summarized by calculating the frequency of significance, average log-fold change, and average adjusted p-value across all iterations.

### Pathway Enrichment

Enrichment analysis was conducted in the Ingenuity Pathway Analysis (IPA, v.23.0, Qiagen) on all genes that had passed the 5% FDR significance criterion and were performed only in those comparisons where the number of DEGs was higher than 20. Pathways were considered significantly enriched if the negative log-transformed Benjamini-Hochberg adjusted p-value (–log(B-H p-value) was greater than 1.5. Based on the known roles of different genes and their relationship in the functionally predefined pathways, IPA calculates a z-score value, which represents the predicted direction of change of the path (activation: z > 0; inhibition: z < 0). For the pathway comparison plot, we selected pathways using two criteria: (1) a high–log₁₀ (BH p-value), and (2) biological relevance to stress resilience and susceptibility, either based on prior studies or known functional involvement in related processes. Top pathways meeting both criteria were then identified and plotted. The analysis included only the pathways associated with the mouse nervous system, neurons, other tissues, and unspecified cell types within the IPA framework.

## Supplementary figures

**Supplementary figure 1.**
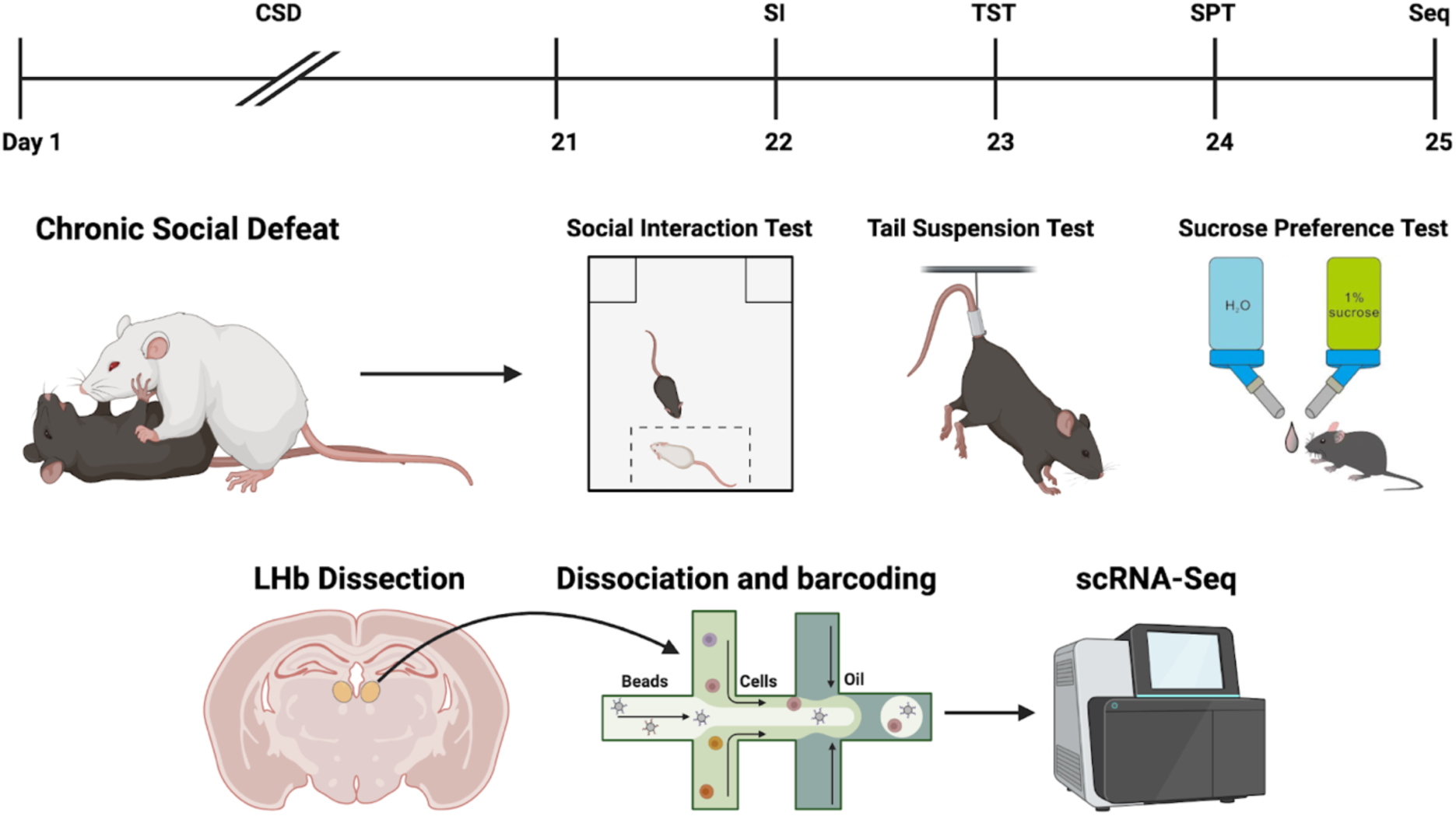
Experimental timeline and methods utilized in the study. Mice were exposed to chronic social defeat stress (CSDS) on 20 consecutive days and were then subjected to the social interaction (SI) test to evaluate social avoidance, tail suspension test (TST) to evaluate despair-like behavior, and the sucrose preference test (SPT) to evaluate anhedonia. Mice with single depressive-like phenotypes, along with stress-resilient and stress-naïve controls, were selected for single-cell RNA-sequencing (scRNA-Seq) of the habenular complex.

**Supplementary figure 2.**
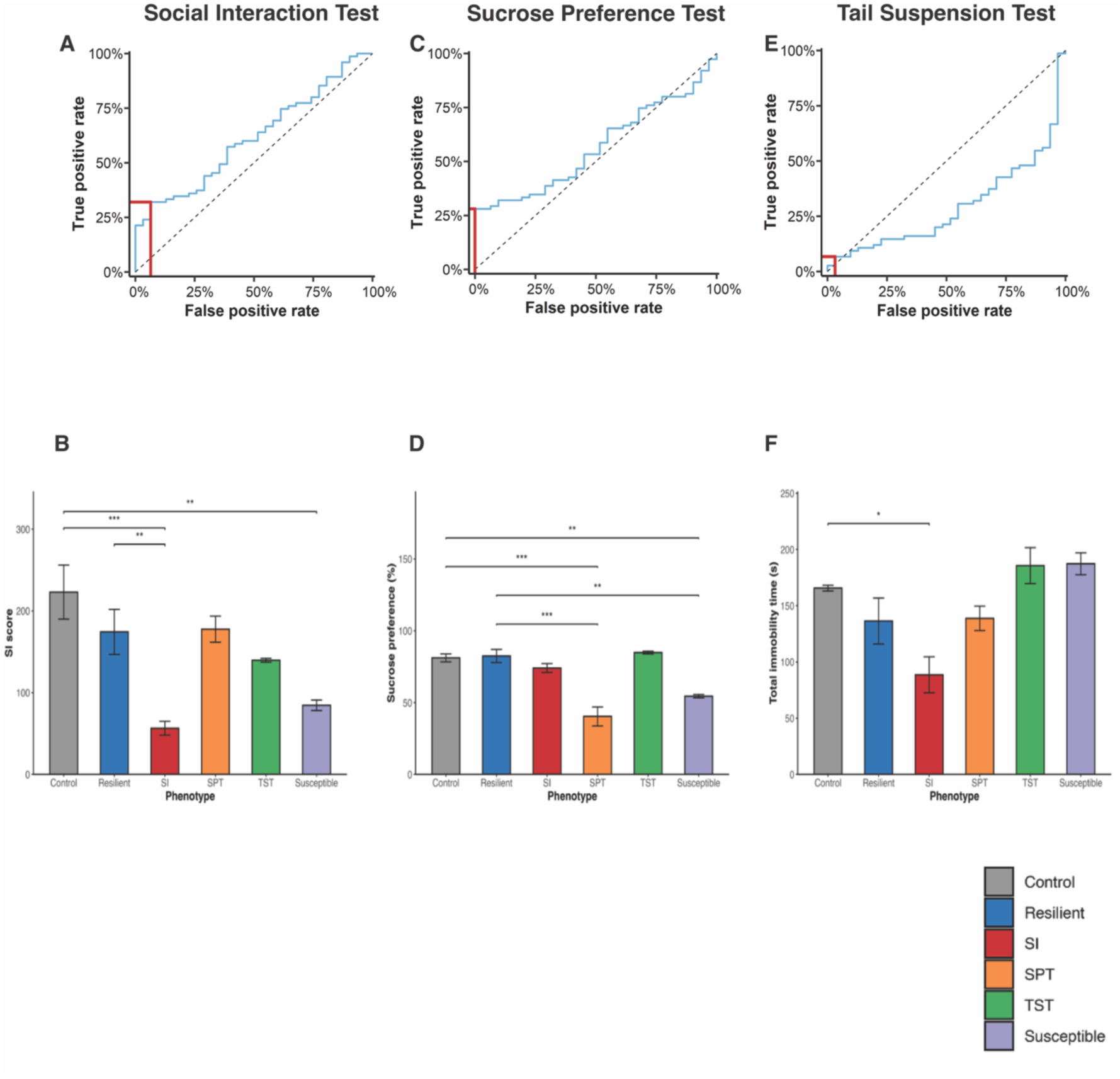
Receiver operating characteristic (ROC) curves of behavioral assays and phenotypic comparisons of behavioral measures. (A,C,E) ROC curves representing the true positive rate (TPR) and false positive rate (FPR) of a binary classifier for different threshold values on the social interaction test (A), sucrose preference test (C), and tail suspension test (E). The dashed line represents the performance of a random classifier, and the red lines highlight the optimal TPR-FRP pairs used to select threshold values for the behavioral measures. (B,D,F) Behavioral measure comparisons between single behavioral phenotypes and stress-resilient and stress-naïve controls for the social interaction test (B), sucrose preference test (D), and tail suspension test (F).Note: *<0.05, **<0.01, ***<0.001

**Supplementary figure 3:**
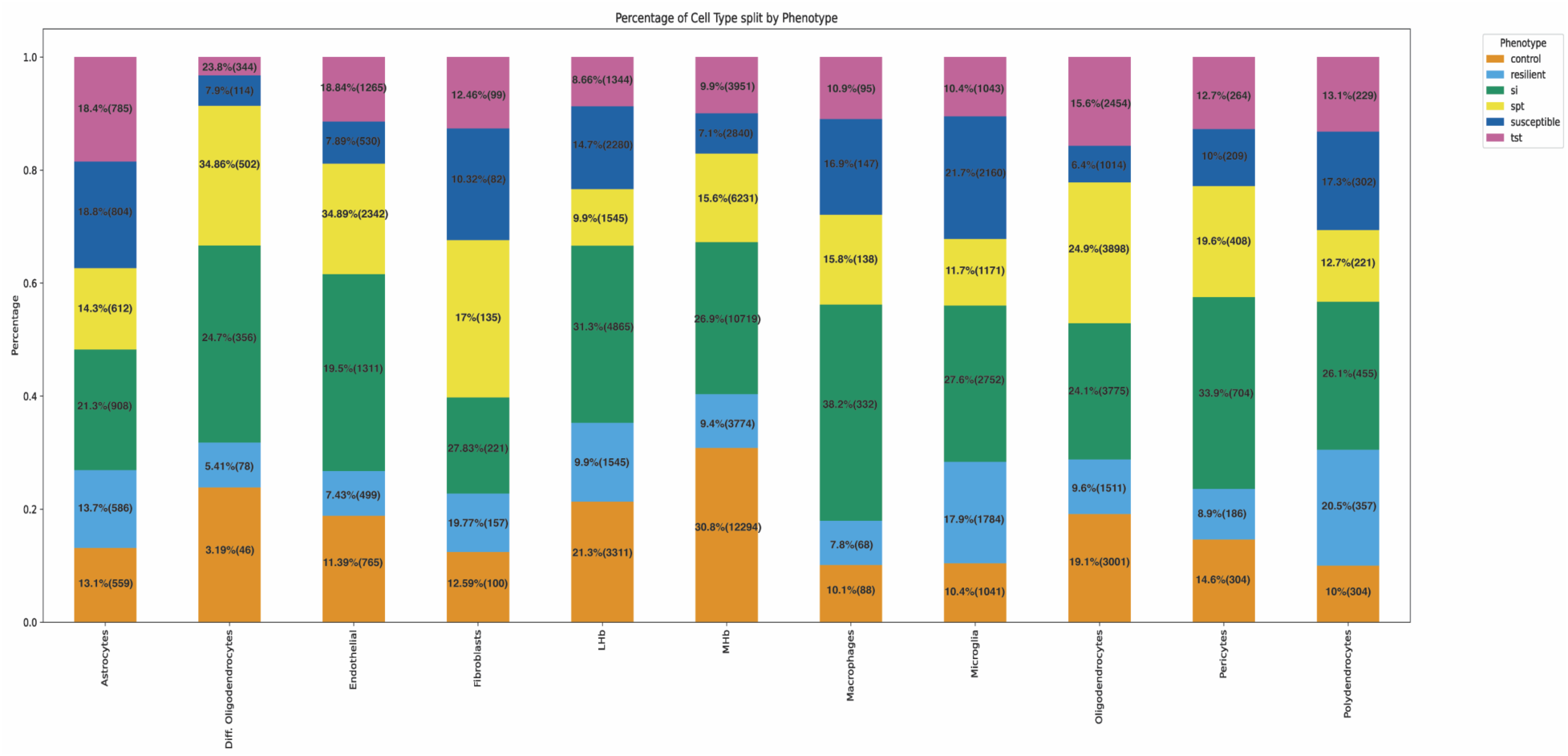
Cell type composition across behavioral phenotypes. Stacked bar plots showing the relative proportions of major cell types within each behavioral phenotype (control, resilient, SI, SPT, susceptible, and TST). This analysis confirmed that transcriptional differences between phenotypes were not confounded by unequal representation of specific cell types.

**Supplementary figure 4:**
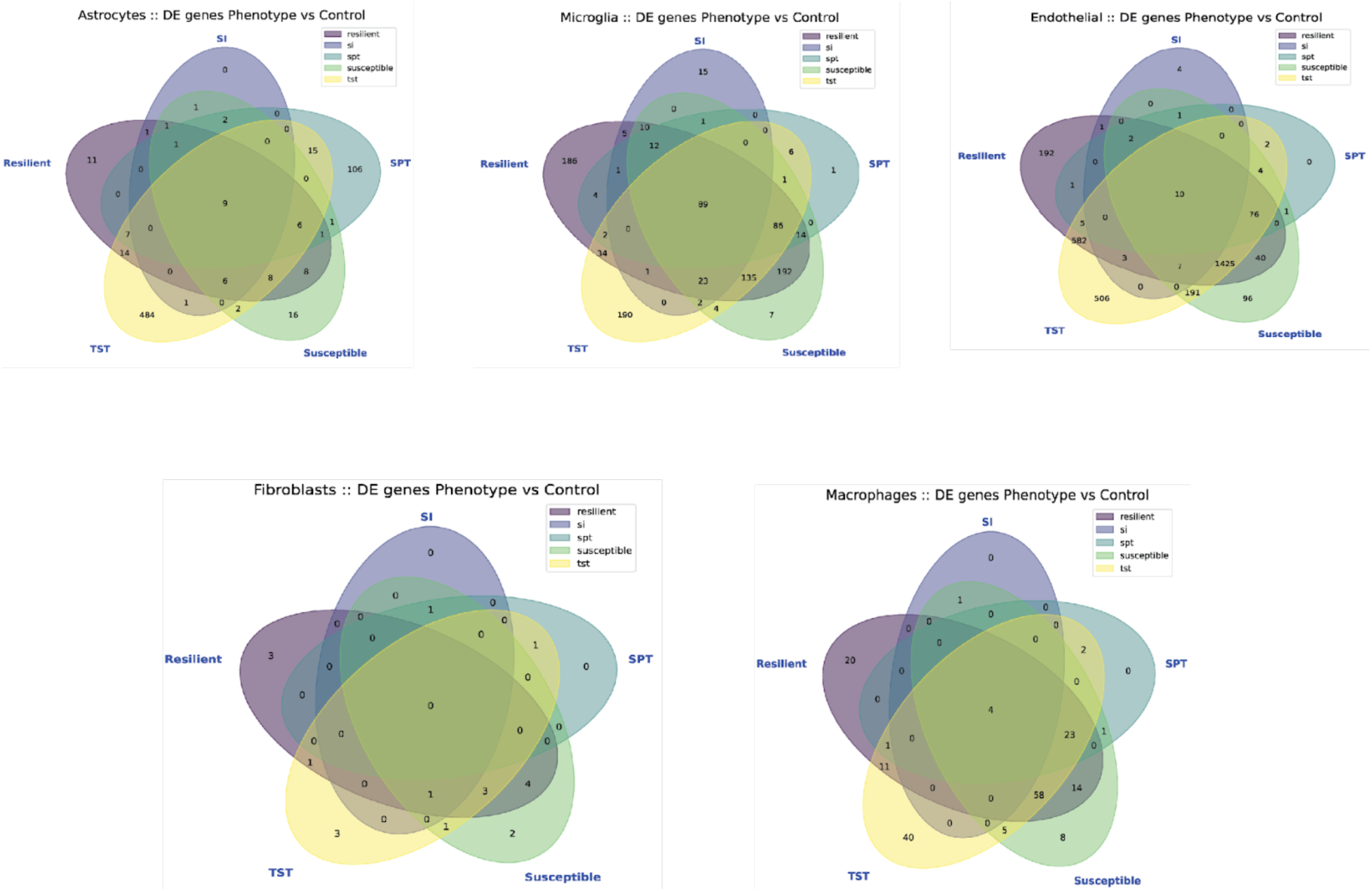
This image displays six Venn diagrams, each representing the DE genes in specific cell types when comparing a Phenotype group against the Control group. The cell types examined are Astrocytes, Microglia, Endothelial cells, Fibroblasts, and Macrophages. Each Venn diagram illustrates the overlap and unique sets of DE genes across four categories: Resilient, SI, SPT, Susceptible, and TST. The numbers within the overlapping regions and unique segments represent the count of DE genes shared or specific to those categories.

**Supplementary figure 5.**
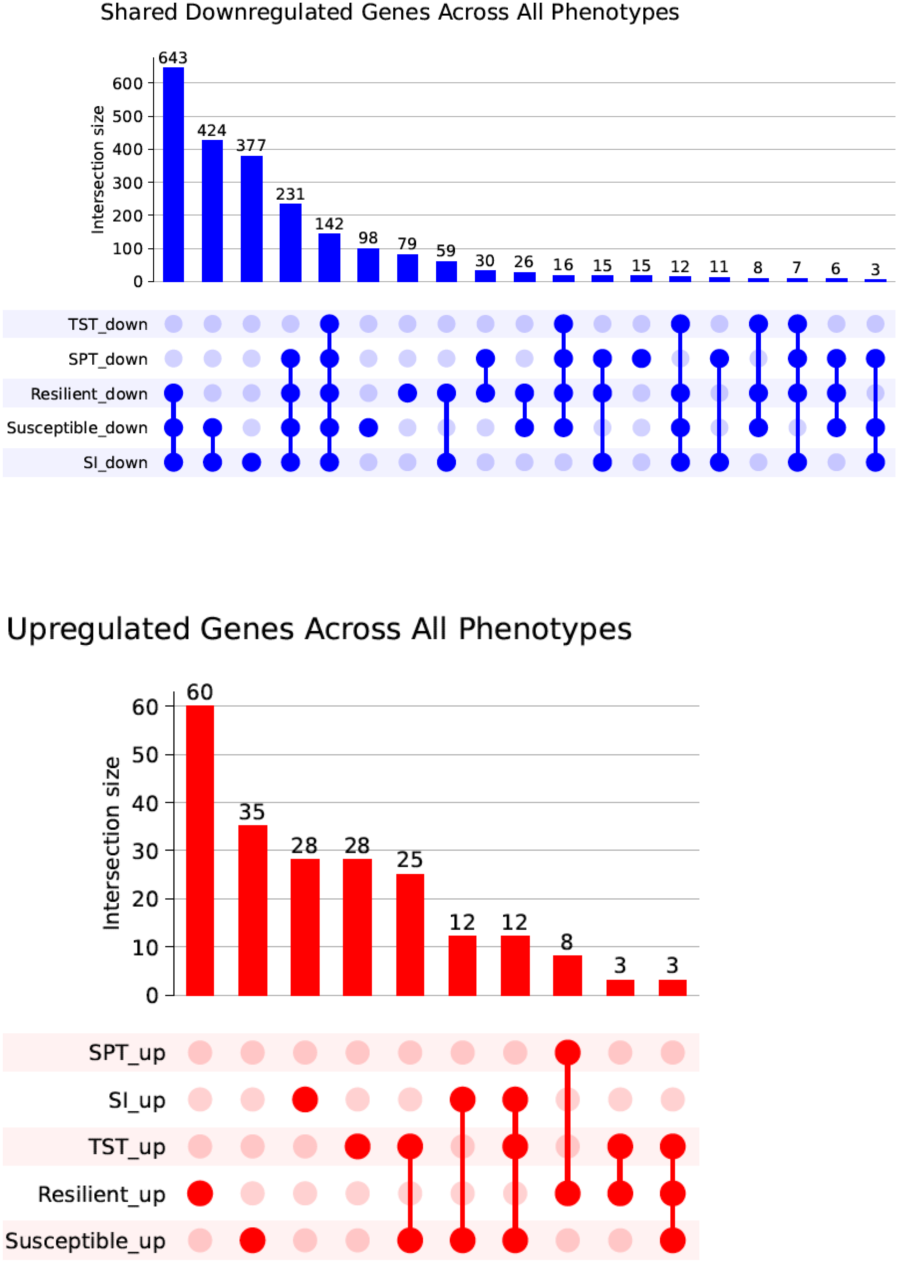
The UpSet plots display intersections of differentially expressed genes across the five phenotypes (SPT, SI, TST, Resilient, Susceptible). Blue color represents downregulated genes, and red color represent upregulated genes. For each plot, bar heights indicate the size of each gene intersection, while the filled and connected dots show which phenotypes contribute to each intersection set.

## References

1. Athira, K. V. et al. An Overview of the Heterogeneity of Major Depressive Disorder: Current Knowledge and Future Prospective. CN 18, 168–187 (2020).

2. Buch, A. M. & Liston, C. Dissecting diagnostic heterogeneity in depression by integrating neuroimaging and genetics. Neuropsychopharmacol. 46, 156–175 (2021).

3. Tang, L. et al. Dissecting biological heterogeneity in major depressive disorder based on neuroimaging subtypes with multi-omics data. Transl Psychiatry 15, 72 (2025).

4. Thorp, J. G. et al. Genetic heterogeneity in self-reported depressive symptoms identified through genetic analyses of the PHQ-9. Psychol. Med. 50, 2385–2396 (2020).

5. Nguyen, T.-D. et al. Genetic heterogeneity and subtypes of major depression. Mol Psychiatry 27, 1667–1675 (2022).

6. Deif, R. & Salama, M. Depression From a Precision Mental Health Perspective: Utilizing Personalized Conceptualizations to Guide Personalized Treatments. Front. Psychiatry 12, 650318 (2021).

7. Fernandes, B. S. et al. The new field of ‘precision psychiatry’. BMC Med 15, 80 (2017).

8. Kambeitz-Ilankovic, L., Koutsouleris, N. & Upthegrove, R. The potential of precision psychiatry: what is in reach? Br J Psychiatry 220, 175–178 (2022).

9. Williams, L. M. Precision psychiatry: a neural circuit taxonomy for depression and anxiety. The Lancet Psychiatry 3, 472–480 (2016).

10. Michel, L., Molina, P. & Mameli, M. The behavioral relevance of a modular organization in the lateral habenula. Neuron 112, 2669–2685 (2024).

11. Piper, J. A., Musumeci, G. & Castorina, A. The Neuroanatomy of the Habenular Complex and Its Role in the Regulation of Affective Behaviors. JFMK 9, 14 (2024).

12. Chaudhury, D. et al. Rapid regulation of depression-related behaviours by control of midbrain dopamine neurons. Nature 493, 532–536 (2013).

13. Fernandez, D. C. et al. Light Affects Mood and Learning through Distinct Retina-Brain Pathways. Cell 175, 71–84.e18 (2018).

14. Liu, Y. et al. Lateral habenula induces cognitive and affective dysfunctions in mice with neuropathic pain via an indirect pathway to the ventral tegmental area. Neuropsychopharmacol. 50, 1039–1050 (2025).

15. Proulx, C. D., Hikosaka, O. & Malinow, R. Reward processing by the lateral habenula in normal and depressive behaviors. Nat Neurosci 17, 1146–1152 (2014).

16. Yang, Y., Wang, H., Hu, J. & Hu, H. Lateral habenula in the pathophysiology of depression. Current Opinion in Neurobiology 48, 90–96 (2018).

17. Ables, J. L., Park, K. & Ibañez–Tallon, I. Understanding the habenula: A major node in circuits regulating emotion and motivation. Pharmacological Research 190, 106734 (2023).

18. Hashikawa, Y. et al. Transcriptional and Spatial Resolution of Cell Types in the Mammalian Habenula. Neuron 106, 743–758.e5 (2020).

19. Wallace, M. L. et al. Anatomical and single-cell transcriptional profiling of the murine habenular complex. eLife 9, e51271 (2020).

20. Matsumoto, M. & Hikosaka, O. Lateral habenula as a source of negative reward signals in dopamine neurons. Nature 447, 1111–1115 (2007).

21. Matsumoto, M. & Hikosaka, O. Two types of dopamine neuron distinctly convey positive and negative motivational signals. Nature 459, 837–841 (2009).

22. Shabel, S. J., Wang, C., Monk, B., Aronson, S. & Malinow, R. Stress transforms lateral habenula reward responses into punishment signals. Proc. Natl. Acad. Sci. U.S.A. 116, 12488–12493 (2019).

23. Cameron, S., Weston-Green, K. & Newell, K. A. The disappointment centre of the brain gets exciting: a systematic review of habenula dysfunction in depression. Transl Psychiatry 14, 499 (2024).

24. Zhukovskaya, A. et al. Heightened lateral habenula activity during stress produces brainwide and behavioral substrates of susceptibility. Neuron 112, 3940–3956.e10 (2024).

25. Chen, M. et al. Brain region–specific action of ketamine as a rapid antidepressant. Science 385, eado7010 (2024).

26. Wang, Z. et al. Deep brain stimulation of habenula reduces depressive symptoms and modulates brain activities in treatment-resistant depression. Nat. Mental Health 2, 1045–1052 (2024).

27. Hu, H., Cui, Y. & Yang, Y. Circuits and functions of the lateral habenula in health and in disease. Nat Rev Neurosci 21, 277–295 (2020).

28. Liu, X. et al. Burst firing in Output-Defined Parallel Habenula Circuit Underlies the Antidepressant Effects of Bright Light Treatment. Advanced Science 11, 2401059 (2024).

29. Zhang, G.-M., Wu, H.-Y., Cui, W.-Q. & Peng, W. Multi-level variations of lateral habenula in depression: A comprehensive review of current evidence. Front. Psychiatry 13, 1043846 (2022).

30. Cerniauskas, I. et al. Chronic Stress Induces Activity, Synaptic, and Transcriptional Remodeling of the Lateral Habenula Associated with Deficits in Motivated Behaviors. Neuron 104, 899–915.e8 (2019).

31. Coffey, K. R., Marx, R. E., Vo, E. K., Nair, S. G. & Neumaier, J. F. Chemogenetic inhibition of lateral habenula projections to the dorsal raphe nucleus reduces passive coping and perseverative reward seeking in rats. Neuropsychopharmacol. 45, 1115–1124 (2020).

32. Luo, L. et al. A cell-type-specific circuit of somatostatin neurons in the habenula encodes antidepressant action in male mice. Nat Commun 16, 3417 (2025).

33. Cui, W. et al. Glial Dysfunction in the Mouse Habenula Causes Depressive-Like Behaviors and Sleep Disturbance. J. Neurosci. 34, 16273–16285 (2014).

34. Cui, Y. et al. Astroglial Kir4.1 in the lateral habenula drives neuronal bursts in depression. Nature 554, 323–327 (2018).

35. Aizawa, H. et al. Potassium Release From the Habenular Astrocytes Induces Depressive-Like Behaviors in Mice. Glia 73, 759–772 (2025).

36. Xin, Q. et al. Neuron-astrocyte coupling in lateral habenula mediates depressive-like behaviors. Cell 188, 3291–3309.e24 (2025).

37. Song, S. et al. Role of phospholipase Cη1 in lateral habenula astrocytes in depressive-like behavior in mice. Exp Mol Med 57, 872–887 (2025).

38. Arjmand, S. et al. The intersection of astrocytes and the endocannabinoid system in the lateral habenula: on the fast-track to novel rapid-acting antidepressants. Mol Psychiatry 27, 3138–3149 (2022).

39. Krishnan, V. et al. Molecular Adaptations Underlying Susceptibility and Resistance to Social Defeat in Brain Reward Regions. Cell 131, 391–404 (2007).

40. Lorsch, Z. S. et al. Computational Analysis of Multidimensional Behavioral Alterations After Chronic Social Defeat Stress. Biological Psychiatry 89, 920–928 (2021).

41. Petković, A. & Chaudhury, D. Encore: Behavioural animal models of stress, depression and mood disorders. Front. Behav. Neurosci. 16, 931964 (2022).

42. Golden, S. A., Covington, H. E., Berton, O. & Russo, S. J. A standardized protocol for repeated social defeat stress in mice. Nat Protoc 6, 1183–1191 (2011).

43. Bonnefil, V. et al. Region-specific myelin differences define behavioral consequences of chronic social defeat stress in mice. eLife 8, e40855 (2019).

44. Liu, J. et al. Impaired adult myelination in the prefrontal cortex of socially isolated mice. Nat Neurosci 15, 1621–1623 (2012).

45. Liu, J., Dietz, K., Hodes, G. E., Russo, S. J. & Casaccia, P. Widespread transcriptional alternations in oligodendrocytes in the adult mouse brain following chronic stress. Developmental Neurobiology 78, 152–162 (2018).

46. Xia, F. et al. Understanding the neural code of stress to control anhedonia. Nature 637, 654–662 (2025).

47. Haynes, S. E. et al. Stress History Modulates CRF Neurons to Establish Resilience. Biological Psychiatry Global Open Science 100656 (2025) doi:10.1016/j.bpsgos.2025.100656.

48. Sinha, R., Lacadie, C. M., Constable, R. T. & Seo, D. Dynamic neural activity during stress signals resilient coping. Proc. Natl. Acad. Sci. U.S.A. 113, 8837–8842 (2016).

49. Tchenio, A., Lecca, S., Valentinova, K. & Mameli, M. Limiting habenular hyperactivity ameliorates maternal separation-driven depressive-like symptoms. Nat Commun 8, 1135 (2017).

50. Seo, J.-S., Zhong, P., Liu, A., Yan, Z. & Greengard, P. Elevation of p11 in lateral habenula mediates depression-like behavior. Mol Psychiatry 23, 1113–1119 (2018).

51. Li, B. et al. Synaptic potentiation onto habenula neurons in the learned helplessness model of depression. Nature 470, 535–539 (2011).

52. Browne, C. A., Hammack, R. & Lucki, I. Dysregulation of the Lateral Habenula in Major Depressive Disorder. Front. Synaptic Neurosci. 10, 46 (2018).

53. Liu, H. et al. Blunted diurnal firing in lateral habenula projections to dorsal raphe nucleus and delayed photoentrainment in stress-susceptible mice. PLoS Biol 19, e3000709 (2021).

54. Mondoloni, S. et al. Serotonin release in the habenula during emotional contagion promotes resilience. Science 385, 1081–1086 (2024).

55. Xie, G. et al. Serotonin modulates glutamatergic transmission to neurons in the lateral habenula. Sci Rep 6, 23798 (2016).

56. Wirtshafter, D., Stratford, T. R. & Shim, I. Placement in a novel environment induces Fos-like immunoreactivity in supramammillary cells projecting to the hippocampus and midbrain. Brain Research 789, 331–334 (1998).

57. Kim, W. & Chung, C. Brain-wide cellular mapping of acute stress-induced activation in male and female mice. The FASEB Journal 35, e22041 (2021).

58. Narain, P. et al. Nighttime-specific differential gene expression in suprachiasmatic nucleus and habenula is associated with resilience to chronic social stress. Transl Psychiatry 14, 407 (2024).

59. Youden, W. J. Index for rating diagnostic tests. Cancer 3, 32–35 (1950).

60. Liu, M.-Y. et al. Sucrose preference test for measurement of stress-induced anhedonia in mice. Nat Protoc 13, 1686–1698 (2018).

61. Wolf, F. A., Angerer, P. & Theis, F. J. SCANPY: large-scale single-cell gene expression data analysis. Genome Biol 19, 15 (2018).

